# Nephrotoxicity of Calcineurin Inhibitors in kidney epithelial cells is independent of Calcineurin and NFAT signaling

**DOI:** 10.1101/2021.07.29.454219

**Authors:** Andrea Karolin, Geneviève Escher, Stefan Rudloff, Daniel Sidler

**Affiliations:** Klinik für Nephrologie und Hypertonie, Inselspital, Bern

**Keywords:** Calcineurin inhibitor (CNI), nephrotoxicity, NFAT-independent, protein kinase inhibitors

## Abstract

**Background:** Calcineurin inhibitors (CNI) such as Cyclosporine A (CsA) and Tacrolimus (FK506) are commonly used after renal transplantation in order to suppress the immune system. In lymphoid cells, CsA acts via the Calcineurin-NFAT axis, whereas in non-lymphoid cells, such as kidney epithelial cells, CsA induces Calcineurin inhibitor toxicity (CNT). Up to date, it is unknown via which off-targets CsA induces CNT in kidney epithelial cells.

**Methods:** In vitro experiments using a surrogate marker to measure CNT induction as well as in vivo studies with acute CNT, were used in order to elucidate the underlying molecular mechanism.

**Results:** Inhibition of the NFAT axis does not show any nephrotoxicity. However, inhibition of p38 and PI3K/Akt Kinases showed induction of nephrotoxicity.

**Conclusions:** These findings show that CsA acts NFAT independent on kidney epithelial cells. Moreover, inhibition of certain protein kinases mimic CsA activity on kidney epithelial cells indicating that p38 and PI3K/Akt kinase pathways might be involved in CNT progression on kidney epithelial cells.

## Introduction

Calcineurin Inhibitors (CNI), e.g., Cyclosporin A (CsA), Tacrolimus (FK506) are pivotal drugs for the prevention of rejection after solid organ transplantation. CNI show a highly interindividual pharmacokinetic profile and have a narrow therapeutic range. Even mild overdosing may lead to substantial acute and chronic side effects, including tumour development, infections, metabolic disturbances, and kidney failure. Calcineurin inhibitors induce nephrotoxicity through (reversible) vasoconstriction in the vas afferent with consecutive relative hypoxia and progressive athero- and arteriolohyalinosis, tubular atrophy and interstitial fibrosis ^1^. Recent register studies have demonstrated that virtually all kidney transplanted patients develop signs of chronic CNT within 10 years after kidney transplantation^2,3^.

In lymphocytes, one major activity of CsA is the inhibition of the Calcineurin-NFAT axis leading to repression of transcriptional programs important for activation, proliferation, and cytokine production. This inhibition primarily contributes to the immune suppressive effect of CsA ^4,5^.

Members of the TNF superfamily have a non-redundant role in the pathogenesis of tissue regeneration and wound healing but are also critically involved in chronic inflammatory and fibrosis ^6–8^. We have recently shown that the TNF superfamily member TWEAK (TNFSF12) is indispensable for the development of CNT in mice ^9^. CsA induces the expression of TWEAK’s receptor, Fn14, in kidney epithelial cells, which facilitates inflammatory and fibrotic signals critical for progressive nephrotoxicity. Deficiency for TWEAK or treatment with TWEAK-neutralizing reagents indeed protected animals from acute CNT lesions ^9–12^. Furthermore, administration of recombinant TWEAK (rTWEAK) induced similar disease as when animals are treated with CsA alone. Interestingly the combination of rTWEAK and CsA induced severe tubulopathy and interstitial inflammation ^9^. The receptor Fn14 is low ubiquitously expressed on epithelial cells and can be induced upon stress and injury stimuli ^13^. Fn14 is the only known receptor to the cytokine TWEAK ^14^. TWEAK is produced by infiltrating and tissue-resident immune cells, largely monocytes and neutrophils ^15–17^.

Currently, it is unclear, how CNI (CsA) interfere with kidney epithelial cells and induce nephrotoxicity. The presented work aims investigating the molecular mechanism in CNT. We hypothesize that CsA elicits its inflammatory and fibrotic activities in renal epithelial cells independent of NFAT, but via inhibition kinase pathways.

## Materials and Methods

### In vitro experiments

The murine kidney epithelial cell line (MCT) has been described previously ^18^. Cells were grown in 10% fetal calf serum (FBS) supplemented Dulbecco modified Eagle medium (DMEM, ThermoFisher) and sub-cultivated in 6- and 12-well format for experiments. Cells were stimulated with various concentrations of either CsA (Sandimmun Neoral® from Novartis) or FK506 (from Cell Signaling, Daveres, MA). In some experiments, cells were incubated with the following reagents: NFAT inhibitor 11R-VIVIT (Merck, 480401), p38 inhibitor SB203580 (Sigma, S8307), p38 inhibitor SB202190 (Sigma, S7076), Gentamycin, Amphotericin B, Cisplatin (Sigma, C2210000) or rTWEAK (peprotec, 310-06). Additionally, in some experiments, HEK293T cells were transfected with various plasmids: pCMV-VIVIT-GFP (Addgene, 11106), pCMV-p38-CA-EGFP and pCMV-eGFP-N1 (Addgene, 6085-1) by using standard Lipofectamine 3000 protocols (ThermoFisher, L3000001).

### Flow cytometry analysis

Flow cytometry was performed on dissociated MCT or HEK293T (when transfected) cells (by Trypsin) stained with Allophycocyanin (APC)-conjugated anti-Fn14 (clone ITEM-4) or respective isotype controls for 30min at 4°C. Proliferation assay was conducted as follows: from naÏve animal’s lymph node/ Spleen T-cells were isolated and stained with 1:1000 e450 proliferation dye (ThermoFisher, 65-0842-85) and treated with various reagents (11R-VIVIT, CsA). Cells were then stimulated with CD3/ CD28 dynabeads for 5 days. For *in vivo* studies, mouse kidneys were digested with 2mg/ml Collagenase I (Sigma, C9891) for 20min at 37°C. Suspension was filtered through a 40μm cell strainer, washed and stained with the following antibodies: Fn14-APC, CD45-PerCP/Cy5-5 (clone: 30-F11), CD11b-APC/Cy7 (clone: M1/70), DAPI, CD326-PE (clone: caa7-9G8). LSR-II Flow Cytometer was used for flow analysis by using the software FACS Diva. The geometric mean fluorescence intensity (MFI) of antibody staining or total count of cells was calculated using FlowJo Software.

### Cell Sort -Aria

Mouse kidney samples were digested and processed as described above. Cells were stained with CD45-PerCP/Cy5.5 and CD326/ EpCAM-PE and DAPI. Cells were sorted according to live cells (DAPI negative). All CD45 positive cells were excluded and only CD326 positive cells were collected and used for RNAseq.

### Kinase Inhibitor Library

MCT were treated with various kinase inhibitors (Enzo, BML-2832) at 10μM. After, cells were dissociated and stained with APC-Fn14. Flow cytometry was used to analyse Fn14 expression on the cell surface.

### Protein Isolation and Analysis

Cells were lysed in Laemmli buffer and separated by sodium dodecyl sulfate polyacrylamide gel electrophoresis and transferred to a polyvinylidene fluoride membrane (ThermoFisher, 88518). Membranes were blocked with 5% milk in TBST for 1h. Incubation with the primary antibody, Fn14 (#4403, Cell Signaling), GAPDH (Merck, cB1001), was conducted overnight at 4°C. The horseradish peroxidase-conjugated secondary antibody was added to the membrane for 1h at RT. Antibodies were diluted according to manufactures recommendation. Visualization of the proteins was conducted using ECL (GE Healthcare, RPN2106) or SuperSignal (ThermoFisher, 34076).

### RNA isolation, reverse transcription and qRT-PCR

Total RNA was isolated from samples using TRIzol reagent (Invitrogen, 15596026). Nanodrop 1000 spectrophotometer was used to measure RNA concentration and evaluate its quality. PrimeScript RT Reagent Kit (Takara, RR037A) was used to generate cDNA. Real-time polymerase chain reaction was performed by using the TaqMan method. For this, probes, which are UPL labelled (Roche) and gene-specific primers were mixed with TaqMan Fast Universal PCR Master Mix (Appliued Biosystems, 4352042). Gene Expression was calculated with the dCt relative to the housekeeping gene. A RT^2^ Profiler PCR Array for mouse nephrotoxicity (PAMM-094Z) was used to determine expression of nephrotoxic genes.

### RNAseq

Total RNA was isolated by using TRIzol reagent according to manufactures protocol. RNA sequencing was performed on illumine NovaSeq 600.

### In vivo experiments: Acute toxicity model

As previously described ^9^, acute CNT lesions were induced by injecting (ip) animals with 100mg/kg CsA (Sandimmun Neoral®), 10mg/kg p38 Inhibitor (SB203580, SB202190) and 10mg 11R-VIVIT on 3 consecutive days twice daily. The fourth day, animals were euthanized, and kidney and blood were collected for subsequent analysis.

## Statistical analysis

Analysis was performed using Prism 5 software (GraphPad, San Diego, CA). Results are the mean ± the standard deviation (SD). Groups were analysed by Student’s t-test (unpaired) or one-way ANOVA. P-values below 0.05 were considered as significant.

## Results

### Surface Fn14 is induced upon CsA exposure in kidney epithelial cells, but not in response to cell-permeable NFAT inhibitors

We have previously demonstrated that the calcineurin inhibitor CsA induces Fn14 surface expression on kidney epithelial cells in vitro and in vivo ^9^. To further support these findings, we challenged MCT cells with various nephrotoxins in a broad range of concentrations. Strikingly, FK506 similarly induced Fn14 induction in MCT cells, while Gentamycin, amphotericin B and Cisplatin did not do so (Figure 1A). As CsA inhibits the NFAT pathway in lymphoid cells, we blunted NFAT activity in kidney epithelial cells by treatment with direct NFAT inhibitors, namely either cell permeable 11R-VIVIT or plasmid-mediated transient expression of VIVIT-plasmid. Prophylactic treatment with 10*μ*M 11R-VIVIT efficiently blocks proliferation of TCRab-stimulated T cells ex vivo (Fig. 1B). Meanwhile, treatment of MCT cells with similar doses of 11R-VIVIT or transient transfection of HEK293T cells with VIVIT-plasmid did not increase Fn14 expression in vitro (Fig. 1C, D). Also, CsA induced further upregulation of Fn14 on cell surfaces even NFAT axis was inhibited (plasmid mediated) (Fig. 1D). It has been shown that CsA sensitizes cells to the cytokine TWEAK^9^. We show in our experiments how cells express significantly higher ICAM (adhesion molecule) after TWEAK treatment when pre-sensitized with CsA. This pre-sensitization is no longer observed when NFAT is inhibited in these cells, comparable to control (Fig. 1E). These experiments indicate, that CsA’s activity in kidney epithelial cells on Fn14 expression cannot be mimicked by direct NFAT inhibitors.

**Figure 1.**
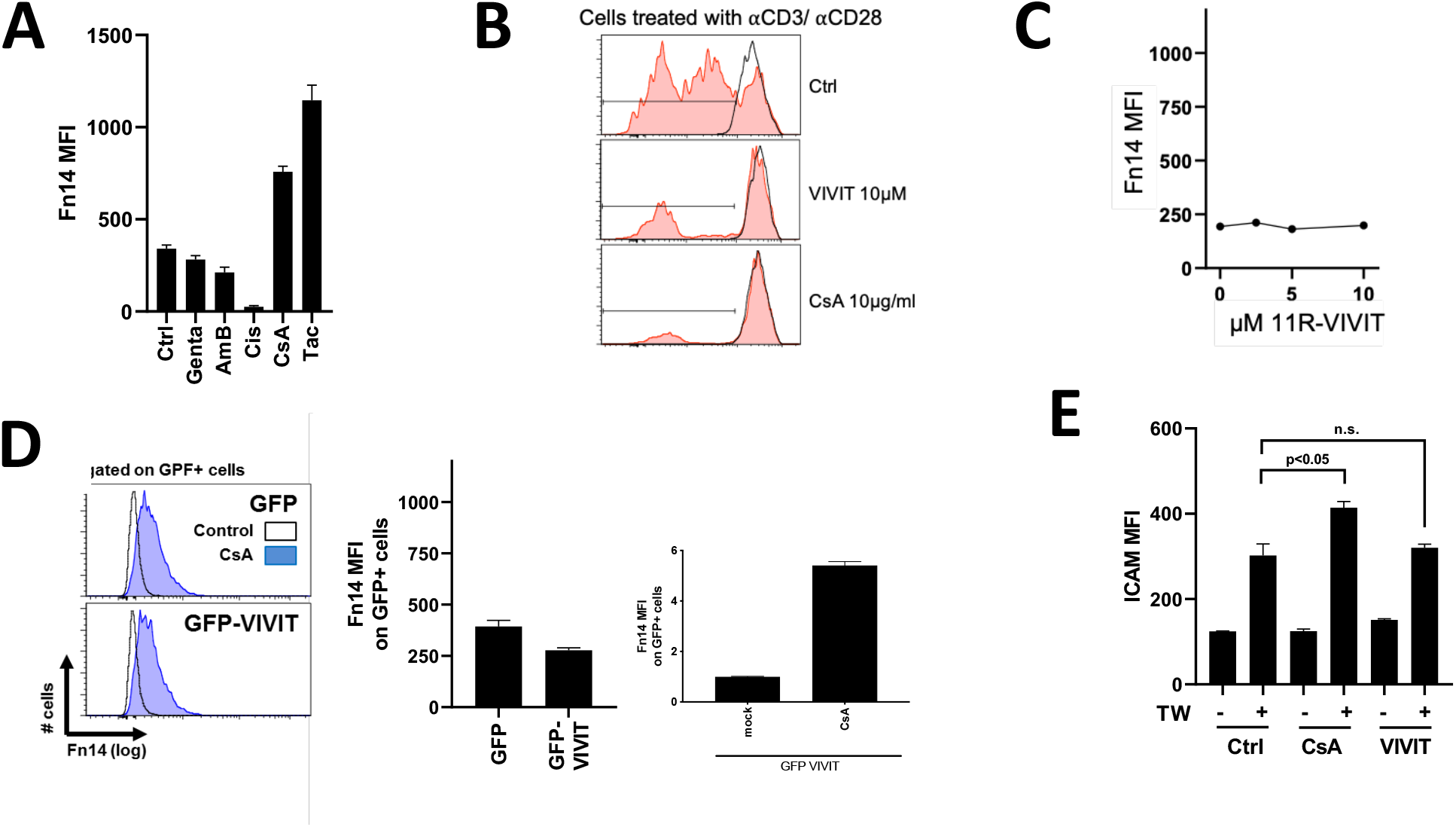
Fn14 induction upon CsA exposure but not upon NFAT inhibition A, Fn14 induction with CsA and Tac (CNIs) but not with Gentamycin, Amphotericin B and Cisplatin. B. NFAT and CsA successfully inhibit proliferation of TCRab-stimulated T cells ex vivo. C-D, inhibition of NFAT axis, by plasmid chemically (11R VIVIT) or (GFP VIVIT), shows no upregulation of Fn14 on cell surface. E, cells sensitized with CsA show upregulation of ICAM upon TWEAK treatment. Inhibition of NFAT combined with TWEAK treatment shows reduced ICAM expression, comparable to control conditions.

### Effects of CsA in kidney epithelial cells are mimicked by inhibitors of protein kinases

Off target effects of pharmacological agents are well known. To investigate CsA’s effect in tubule epithelial cells of the kidneys, we treated MCT cells with the Kinase Inhibitor library (Enzo, BML-2832) and investigated the effect of individual compounds on Fn14 induction as surrogate marker for pro-fibrotic responses. Indeed, only certain protein kinase inhibitors shown in Fig. 2A and 2B significant upregulated Fn14 on the surface of MCT cells. A multitude of factors, such as BAY 11-7082, SB-203580, Wortmannin, GF 109203X, Palmitoyl-DL-carnitine, Triciribine, SB-202190 and Hypericin led to significant induction of surface Fn14. Notably, two p38 kinase inhibitors (SB203580 and SB202190) showed potential to upregulate Fn14 on cell surfaces. Titration of both inhibitors revealed similar potential to induce Fn14 (Fig. 2C). Protein from SB203580 treated cells showed similar upregulation of Fn14 as cells treated with CsA (Fig. 2D). Cells treated simultaneously with a p38 kinase inhibitor and CsA showed an increased expression of Fn14 compared to cells only treated with CsA or one of the p38 kinase inhibitor (Fig. 2E). To corroborate these findings, we generated cells expressing a constitutive active p38 kinase (pCMV-p38-CA-EGFP). Basal Fn14 expression (MFI) was moderate in all untreated mock-transfected (pCMV-eGFP-N1) cells and p38-CA-expressing cells. Mock-transfected control (GFP) showed a highly significant upregulation of Fn14 upon SB203580 or CsA stimulation. Meanwhile, for the p38-CA-expressing cells, the p38 inhibitor or CsA induced significantly lower Fn14 expression (Fig. 2F). Furthermore, a nephrotoxic gene array revealed that CsA upregulates numerous genes involved in toxicity in the kidney. What is more, that SB203580 also shows a high induction potential of nephrotoxic genes (Fig. 2G).

**Figure 2.**
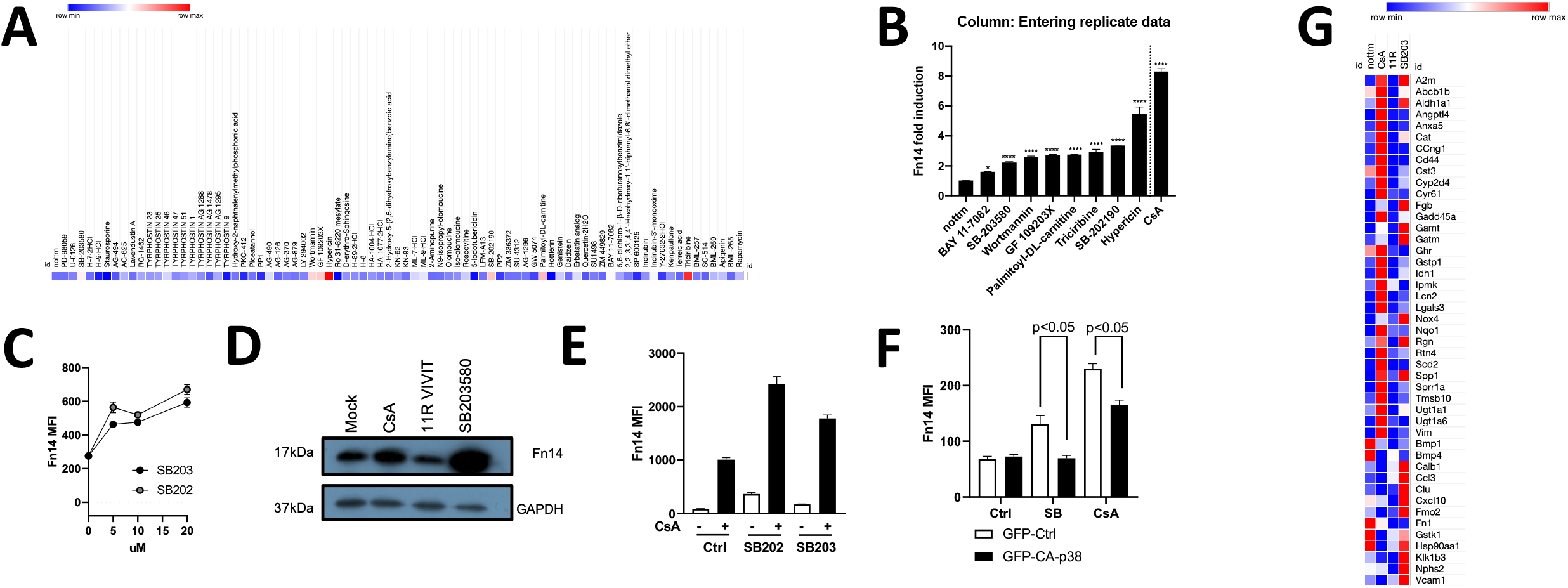
Effects of CsA on kidney epithelial cells are mimicked by protien kinases inhibitors. A-B, screen of a protein kinase library revealed that inhibition of PI3K, Akt, IKK and p38 kinase induces Fn14. C, titration of two p38 inhibitors (SB203580 and SB202190) revealed similar potential to induce Fn14. D, Protein levels of Fn14 are increased upon CsA and SB203580 treatment, but not when NFAT is inhibited. E, increased expression of Fn14 is detected when cells are treated simultaneously with p38 kinase inhibitor and CsA compared to single treatment with inhibitors. F, constitutive active p38 kinase shows reduced Fn14 expression on cells when treated with either SB203580 or CsA. G, nephrotoxic genes are upregulated with CsA as well as with SB203580.

### CsA, but not VIVIT induces nephrotoxicity in vivo

Exposure of mice with high dose of CsA induces nephrotoxicity, reflected in induction of acute phase, fibrosis and inflammatory genes, e.g. Angptl4, CD24 and TNFRSF12a. Furthermore, mice treated with p38 inhibitors SB203580 and SB202190 show similar upregulation of genes involved in nephrotoxicity (Fig. 3A). Meanwhile, animals treated with VIVIT showed no such signatures. Moreover, immune cell infiltration (CD45+) into the kidney is upregulated in animals treated with CsA or p38 kinase inhibitors (Fig. 3B). Not only, could we detect such changes in kidney infiltrated cells but also immune cell compartment, notably CD45+ and CD11b+ cells, seem to be increased upon CNI or p38 kinase treatment (Fig. 3C).

**Figure 3.**
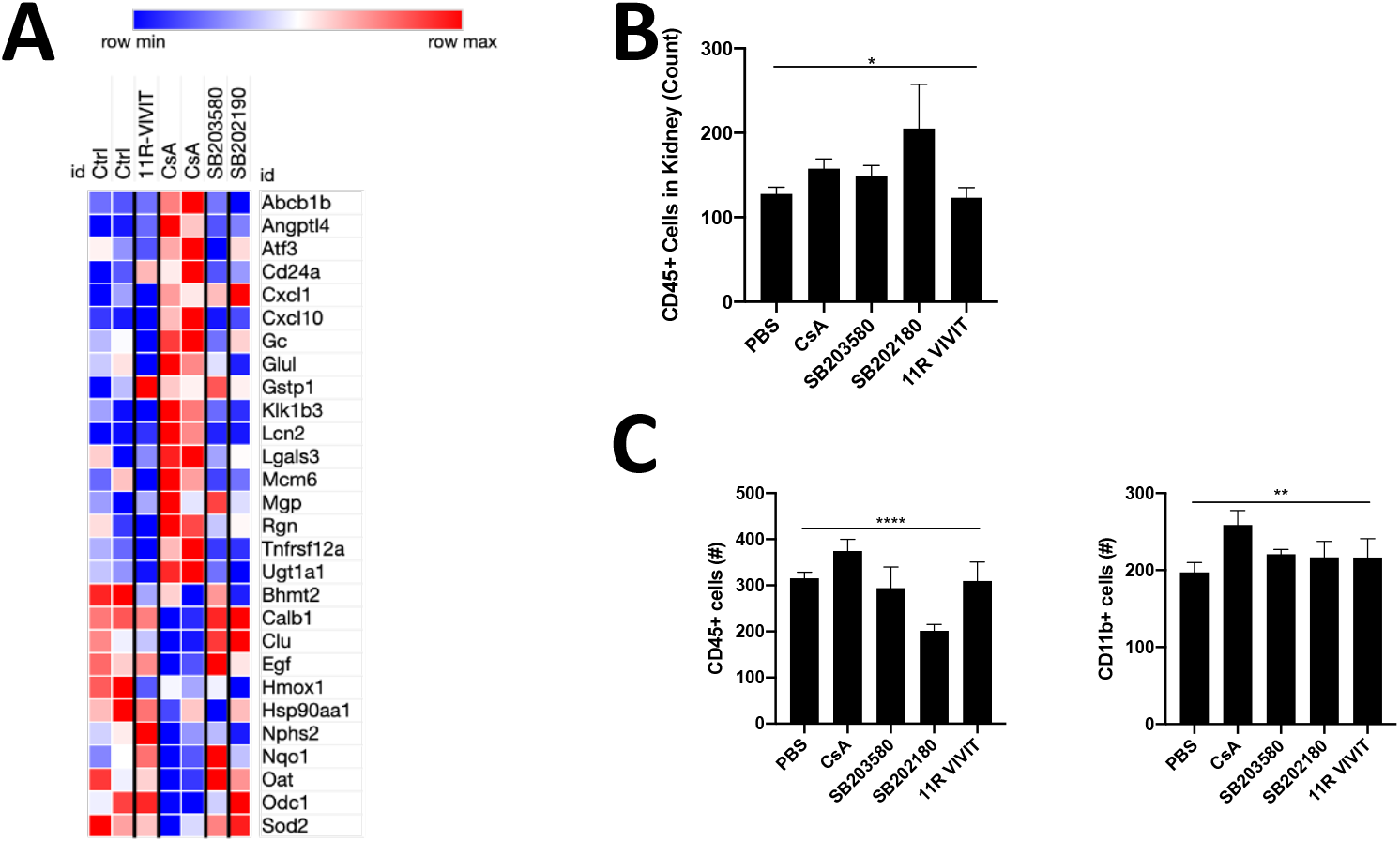
In vivo experiments show development of nephrotoxicity upon CsA and p38-kinase inhibitor. A. nephrotoxic gene array reveals clear upregulation of genes involved in nephrotoxicity upon CsA treatment. Also, both p38 inhibitors revealed upregulation of nephrotoxic genes. B, higher CD45+ infiltrated cells in the kidneys of CsA, SB203580 and SB202190 were detected. C, monocytic and lymphocytic cell compartment in the blood show. A slight increase in numnber in animals treated with CsA and p38-inhibitors.

## Discussions

CNI play a crucial role in effective immunosuppression after kidney transplantation. However, long term treatment lead to chronic nephrotoxicity. In lymphoid cells, CsA and FK506 interfere with the cytosolic phosphatase Calcineurin and thereby prevent dephosphorylation of nuclear factor of activated T cells (NFAT) and consequently gene transcription. To date, it is unclear, if biological effects of CNI in non-lymphoid cells result similarly from disrupted Calcineurin-NFAT signalling. This work aimed to investigate the molecular mechanism of CNT in epithelial cells. We present experimental evidence, that CNT lesions are induced in kidney epithelial cells in vivo and in vitro independent of canonical Calcineurin – NFAT signalling, yet induced by kinase inhibitors, including inhibitors of the p38, IKK and PI3K pathway.

Firstly, we showed that Fn14 upregulation is achieved by treatment of kidney epithelial cells by CNI (CsA and FK506) whereas other compounds fail to upregulate Fn14. Furthermore, we show that also an effective inhibitor (11R-VIVIT) of the NFAT axis fails to induce Fn14 in epithelial cells. These results confirm previous studies ^9^ that Fn14 is specifically upregulated in kidney epithelial cells when treated with a nephrotoxic agent and that Fn14 is a good surrogate marker to measure nephrotoxicity and pro-fibrotic signals. Furthermore, did our result show that nephrotoxic CNI signalling in kidney epithelial cells is not mediated via the canonical Calcineurin-NFAT pathway in vitro and in vivo.

Likely, CNT is the consequence of off-target effects of CsA in kidney epithelial cells. It has been hypothesized that CsA targets disparate signalling pathways in lymphoid and non-lymphoid cells ^19^ . As we showed that CsA’s fibrotic effect on kidney cells is stable and easy to measure (Fn14) and that the nephrotoxic effect is NFAT-independent, we show here synthetic compounds and their capacity to induce Fn14. Our screen reveals that protein inhibitors of the p38, PI3K/Akt and PKC kinases mimic CsA’s activity on epithelial cells. These effects are specific for respective pathways, since chemically unrelated inhibitors of these routes elicited similar activities (SB203580 and SB202580/ Wortmannin and Triciribine/ Hypericin, GF 109203X and Palmitoyl-DL-Carnitine). Meanwhile, inhibitors of the MEK1/2-ERK (PD-98059/ U-0126) and JNK pathway (SP600125) had no Fn14-inducing activities. Results strongly suggest that the CNT of CsA on kidney epithelial cells acts by inhibition of p38 MAPK and PI3K/Akt kinases. Both kinase pathways, p38 and PI3K/Akt are important in epithelial cells for differentiation, proliferation and apoptosis and extracellular matrix synthesis. In both pathways the GSK3β involved^20^. Our top-down approach reveals that the inhibition of GSK3β (by Indirubin-3’-monooxime) shows slight upregulation of Fn14.

This work has several limitations. First, induction of sustained CNT lesions is cumbersome in inbred mice and treatments require high to supramaximal CsA doses. The reason for this notion is unknown, yet likely lies in a greater renal reserve for hemodynamic toxicity, low-renin levels, and other factors ^3,21,22^. Possibly, single nucleotide polymorphisms further contribute to this CsA resistance in inbred mice. CNT lesions are not highly specific to CsA or Tac exposure in mice and humans. Similar lesions, including arteriohyalinosis and interstitial fibrosis/tubular atrophy (IFTA) can be identified in patients with chronic active and chronic inactive rejection and in kidney transplantation from older and hypertensive donors ^23,24^.

This work offers potential for future experimental and clinical projects. First, numerous nephroprotective treatments are available or awaiting clinical approval, including SGLT-2 inhibitors ^25^, PLG1-analoga ^26^ and Finerenone ^27^. Since CNT is likely an off-target effect of CsA directly on kidney epithelial cells, such nephroprotective drugs could be of great value to prevent/treat developing or existing CNT in solid organ transplant recipients.

CNT shows a highly variable clinical presentation, and some patients develop early and progressive lesions within months of SOT, while others tolerate similar CNT doses/through levels over many years or decades without significant graft or kidney function impairment and absence of CNT lesions in biopsy specimen ^3,28,29^. The reason for this highly inter-individual predisposition is unclear, but likely lies in single nucleotide polymorphisms (from the donor in a setting of kidney transplantation, and the recipient). To date, no GWAS study has been performed to decipher risk genes for rapid and progressive CNT after SOT.

In conclusion, these results suggest that CsA acts NFAT-independently on kidney cells yet requires p38 kinase and PI3K/Akt kinase pathways. Furthermore, could we show in vitro and in vivo that inhibition of proposed pathways (p38 and PI3K/Akt) mimics the toxicity of CsA on epithelial cells. However, we cannot exclude that CNT is rather not a highly specific interaction with a single molecular pathway.

## Acknowledgment

We thank Michael Croft, La Jolla Institute for Allergy and Immunology, California, for his helpful inputs and discussions.

## Notes

Conflict of Interest All authors deny any conflict of interest that might bias their work.

### Competing Interest Statement

The authors have declared no competing interest.

